# NMDA receptor-dependent Hebbian plasticity refines hippocampal spatial representations during two-dimensional navigation learning

**DOI:** 10.64898/2026.07.13.734058

**Authors:** Ronen Reshef, Mina Shahi, Victoria Ho, Matthias Ollivier, Ahmet Arac, Ariel Cohen, David Yamin, Amanda Tran, Ruth Tjondropurnomo, Baljit S. Khakh, Daniel Aharoni, Thomas J. O’Dell, Peyman Golshani

**Affiliations:** Department of Neurology, David Geffen School of Medicine, University of California, Los Angeles, Los Angeles, CA, USA; Department of Physiology, David Geffen School of Medicine, University of California, Los Angeles, Los Angeles, CA, USA; Greater Los Angeles VA Medical Center, Los Angeles, CA, USA; Semel Institute of Neuroscience and Human Behavior, David Geffen School of Medicine, University of California, Los Angeles, Los Angeles, CA, USA; Intellectual and Developmental Disability Research Center, David Geffen School of Medicine, University of California, Los Angeles, Los Angeles, CA, USA; Integrative Center for Learning and Memory, David Geffen School of Medicine, University of California, Los Angeles, Los Angeles, CA, USA

## Abstract

Hippocampal place cell activity represents an animal’s location in space; yet, how hippocampal neuronal population dynamics change with spatial learning and the mechanisms underlying these activity changes, which drive allocentric navigation to a learned goal, are poorly understood. To address these questions, we performed calcium imaging with a novel wire-free waterproof miniaturized microscope to image the activity of large populations of hippocampal CA1 neurons during spatial learning of a two-dimensional navigational task, the Morris water maze. We followed the same cells during learning and were able to directly examine how each neuron in the ensemble, and the ensemble as a whole, changes its response properties. We found that neuronal spatial selectivity increased and population decoding of spatial location improved as mice learned to navigate to the goal. Viral CRISPR knock out of *Grin1* (encoding the essential GluN1 NMDA receptor subunit) in dorsal hippocampal neurons, dramatically reduced long-term potentiation in CA1. This manipulation also prevented the increase in spatial selectivity and improvement of population decoding with spatial learning and resulted in learning deficits in the Morris water maze. Together, our results show that dorsal hippocampus NMDAR-dependent synaptic plasticity is essential for the learning-dependent refinement of CA1 place selectivity and improvement in population decoding of space.

## Introduction

To survive, animals must use distal cues to navigate the best route to specific remembered locations in space. Navigation using distal cues has been hypothesized to be based on the representation of spatial locations in hippocampal CA1 neurons termed place cells, which collectively form a cognitive map of the environment (*1*). Several studies have shown that the dorsal hippocampal region is critical for learning and performance of a goal-directed 2-dimensional (2D) spatial navigation task using distal cues (*2*, *3*). However, whether CA1 neuronal place representations are shaped by learning and the mechanisms underlying these changes that drive allocentric navigation to a goal in a 2D environment are poorly understood.

Several studies have shown that spatial learning results in an accumulation or reorganization of place fields near the goal (*4–8*). This reorganization may mark the goal (*9*), but in 2D navigation, it cannot explain how animals navigate using direct paths to the goal when they are remote from this goal and not activating the added place fields. Other studies have found improved precision of coding for place in individual cells during learning in one-dimensional environments or during 2D random foraging, both of which impose minimal navigational demands (*10–13*). Although N-methyl-D-aspartate receptor (NMDAR) dependent synaptic plasticity was hypothesized as a plausible mechanism mediating learning-dependent changes in place cell precision (*14*, *15*) and deletion of this receptor in the hippocampus was shown to degrade spatial representations (*16*–*18*), these findings were obtained in constrained non-navigation-directed settings.

One study employing virtual reality navigation in 2D found evidence of distance coding and path integration in CA1, rather than alterations in spatial coding after spatial learning (*19*), offering an alternative explanation for the neural dynamics underlying learning. Yet, given that navigation in virtual reality dramatically alters place coding in 2D (*20*), this approach may force animals to rely on alternative strategies.

It has remained a major challenge to follow the representations of the same neurons during learning in a well-controlled 2D freely behaving spatial navigation task, and to probe the mechanisms underlying spatial learning by causally manipulating plasticity in the same population. To meet this challenge, we examined how place cell representations in the same neurons are shaped by spatial learning across days in a 2D allocentric navigational task. We found that hippocampal spatial representations become more precise after learning the Morris water maze, and that these changes are prevented by CRISPR deletion of NMDA receptors in the dorsal hippocampus, a manipulation that blocked long-term potentiation and prevented learning.

## Results

To study how spatial representations change during spatial learning, we monitored hippocampal neural activity by performing calcium imaging with wire-free miniaturized microscopes (*21*, *22*). We chose to image CA1 activity while animals performed the Morris water maze (MWM) hidden platform test, because in this task animals are forced to use a single navigational strategy by using distal visual cues to locate a goal in a 2D arena (*23*). Furthermore, the task has been highly validated in the learning and memory literature and proven to be hippocampal-dependent (*24*).

To image in the MWM, we used wire-free water-proof miniaturized microscopes and minimized the weight load by attaching each miniscope to three helium balloons (Fig. 1, A; see Methods). We imaged animals as they performed a well-established behavioral protocol; animals performed 3 blocks of behavior, with each block consisting of 4 consecutive trials, completing 12 trials daily. This protocol resulted in increased imaging time compared to other variants of the MWM with fewer trials, allowing us to better probe neural mechanisms of spatial learning (*25*) (see Methods). Over 6 days of training, mice displayed a significant progressive reduction in the latency to locate the platform (Fig. 1, B). Search trajectories varied until reaching maximum performance on Day 6 (Supplement Fig. 1, A). Because animals found the platform more rapidly on Day 6 resulting in reduced imaging time, we added another 12-24 trials on Day 6 after the probe trial to equalize the time imaged. On Day 1, animals searched for the maze relatively uniformly, while on Day 6 animals took more direct paths to the goal (Fig. 1, C). Probe memory test trials performed on Day 3 and Day 6, showed that there was a significant increase in exploration of the target quadrant on Day 6 but not on Day 3, demonstrating that animals had learned the location of the platform by Day 6 (Fig. 1, D-E; Supplement Fig. 1, B-C).

**Fig 1:**
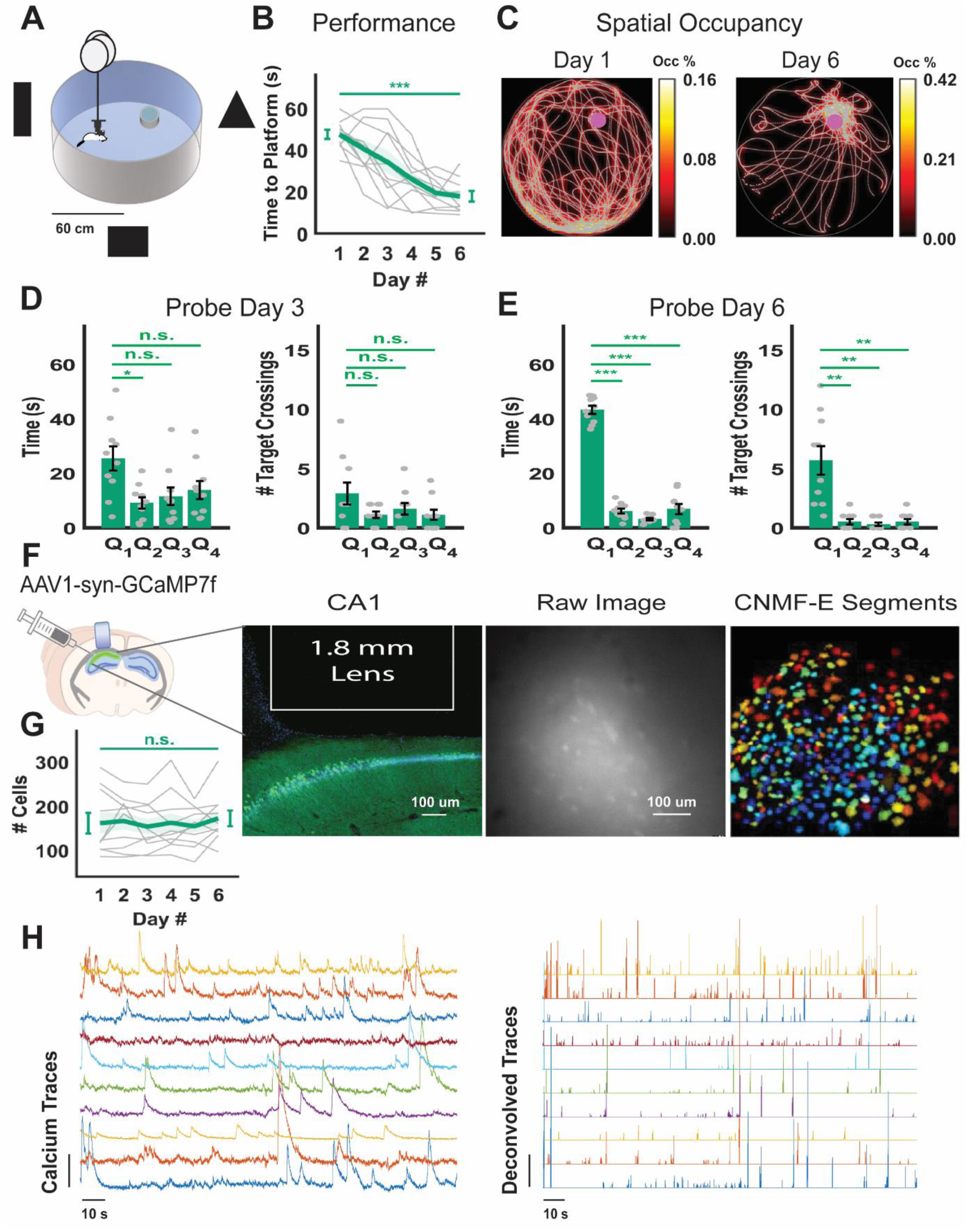
Imaging the CA1 neuronal population during spatial learning in the MWM. (a) Schematic of experimental setup. (b) Progression of learning in the MWM across six days of training in the MWM, as measured by latency to reach the platform on each trial, averaged for each day (n= 10 mice; *** P≤ 0.001, paired *t*-test). (c) Spatial occupancy in the maze across all trials of Day 1 and Day 6, measured as percentage of time spent in each maze location for each imaging day. (d) Spatial memory performance in the probe trial on Day 3, quantified as the proportion of time spent in the quadrant of the learned platform location – Q1 (n=10 mice; * P≤ 0.0167; P= 0.08; P= 0.115) and the number of crossings of the platform location – Q1 (n= 10 mice; P= 0.0642; P= 0.281; P= 0.167). (e) Spatial memory performance in the probe of Day 6, quantified as the proportion of time spent in the quadrant of the learned platform location – Q1 (n= 10 mice; *** P≤ 0.001) and the number of crossings of the platform location – Q1 (n= 10 mice; ** P≤ 0.01). (f) Left: schematic of the virus injection and location of lens implantation above the CA1 region of the hippocampus. Histological section showing the aspiration cavity of the GRIN lens and CA1 cells expressing GCaMP7f (green). DAPI nuclear counterstain shown in blue. Middle: raw GCAMP7f fluorescence of the field of view from an example animal. Right: single neurons segmented from the field of view by CNMF-E. (g) The number of neurons imaged across training in the MWM (n= 10 mice; P= 0.576; paired *t*-test). (h) Example of calcium traces from 10 neurons and deconvolved spikes extracted from them using the Oasis package. Statistical test in Figures 1d and 1e was performed with planned paired *t*-tests with Bonferroni correction.

### Improvement of place tuning in CA1 neurons after spatial learning

We imaged neural activity in CA1 during the task acquisition by injecting mice with AAV1-syn-GCaMP7f into CA1 and implanting a GRIN lens directly above the stratum oriens (Fig. 1, F). To measure learning-related changes in spatial tuning of CA1 neurons, we used the binning method of summing the activity in each spatial bin divided by the occupancy in that bin (*2c*). We focused on the activity recorded on Day 1 before learning and Day 6 after learning. Around 1,800 neurons were imaged throughout the training days from 10 animals (Fig. 1, G). We used the deconvolved calcium traces of each neuron to study its firing properties (Fig.1, H).

To understand how tuning changes in the same population of neurons before and after learning, we mainly analyzed neurons that were active both on Day 1 and Day 6. We registered these neurons across days using the CellReg algorithm (*27*). 680 neurons recorded across 10 mice could be registered across Day 1 and Day 6 (Fig. 2, A). To quantify the precision of spatial tuning in these neurons, we calculated a commonly used spatial sparsity measure that examines how selectively each neuron fires across specific positions in the arena (*28*, *29*) (see Methods). We chose sparsity to quantify spatial precision tuning because it is less sensitive to the limited sampling bias (*30–32*). Some neurons fired in a non-selective manner throughout the arena on Day 1, thereby exhibiting low spatial sparsity. However, on Day 6, the same cells showed more spatially selective firing in different locations of the maze, exhibiting increased spatial sparsity (Fig. 2, B). To ensure that differences in behavioral sampling between Day 1 and Day 6 do not confound these results (Fig 2, B; Fig. 1, C), we circularly shuffled the firing of neurons and calculated the sparsity of shuffled firing for each neuron (see Methods). To control for the changes in behavioral sampling, we subtracted the 95^th^ percentile of the sparsity measured for shuffled trials from those recorded during exploration for each neuron. The cumulative distributions of the shuffle-subtracted sparsity values of all animals were statistically different on Day 1 and Day 6, with sparsity increasing on Day 6 (Fig. 2, C). The sparsity was also greater when we averaged the population shuffle-subtracted sparsity scores for each animal and compared the average of all animals from Day 1 to Day 6. The average shuffle-subtracted sparsity scores on each day became negative because 95^th^ percentile of the shuffle is higher than most cells’ sparsity scores; however, importantly, the rise in the sparsity averages exhibits the learning-dependent changes (Fig. 2. C). The calcium event frequency was similar on Day 6 compared to Day 1 (Supplement Fig. 2, A). The speed of the animals, a known modulator of hippocampal activity (*33*), was not different on these two imaging days (Supplement Fig. 2, B). Results were similar if all neurons imaged were studied, rather than neurons that were only active on both Day 1 and Day 6 (Fig. 2, D). To more carefully control the behavioral bias and to equalize the number of occupied bins, we subsampled Day 1 behavior to match that of Day 6. Using only the same positions (bins) that were visited on both days, as well as using the same amount of data across the days as before, we repeated the analysis. Consistent with previous results, we found the 95^th^ percentile shuffle-subtracted sparsity rising on Day 6 compared to Day 1 in the shared cells (Supplement Fig. 2, C-D), and in all the cells imaged (Supplement Fig. 2, E-F). Therefore, the precision of spatial firing of individual cells increases in CA1 with learning on the MWM.

**Fig 2:**
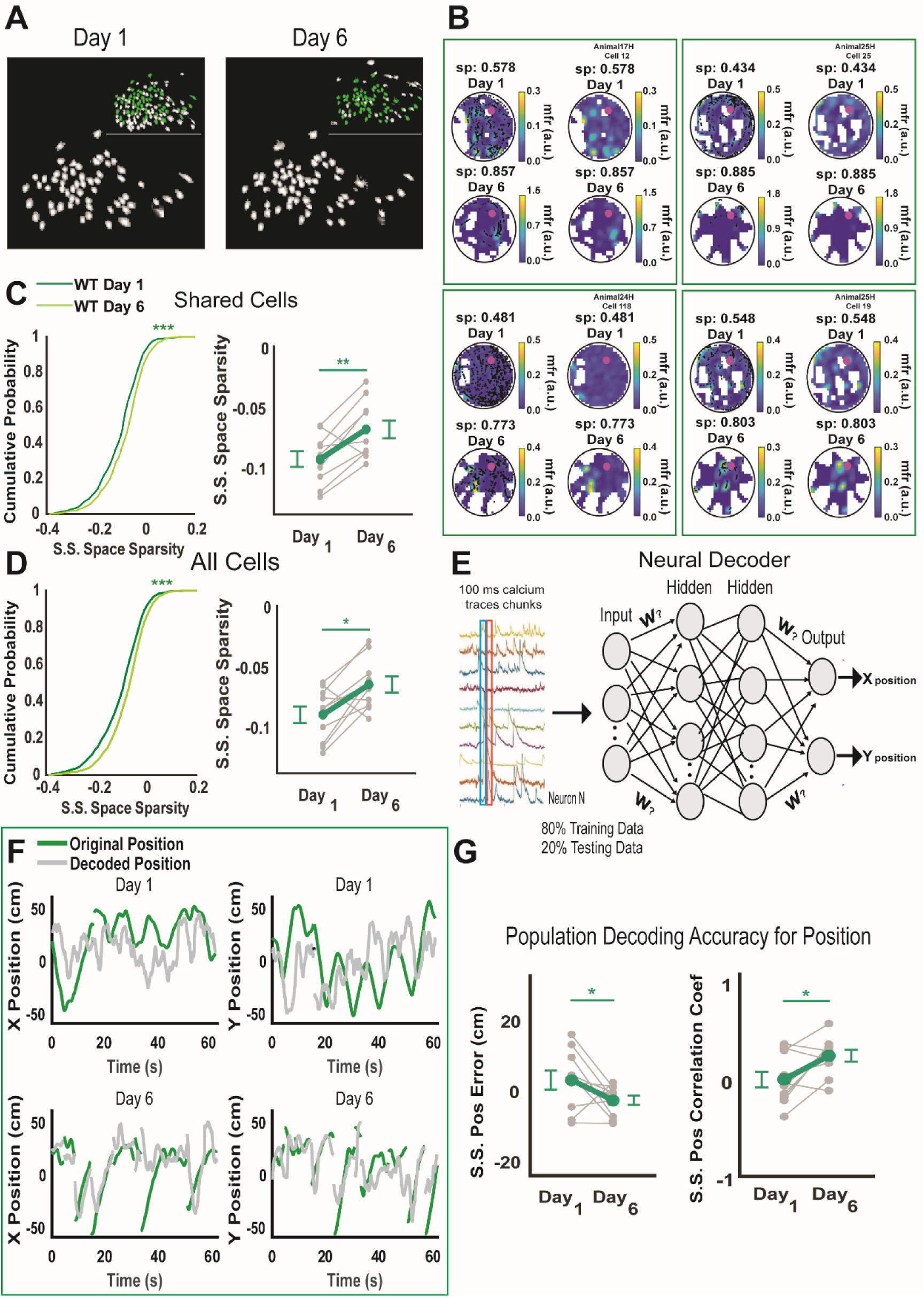
Population of CA1 neurons increases selectivity for place with spatial learning in the MWM. (a) Example of neuron footprints registered as being the same neurons by CellReg on Day 1 and Day 6. Insets: All neurons in white and registered neurons in green. (b) Left column: Heat map of the activity of the imaged neurons on Day 1 and Day 6 with deconvolved events color coded in black. Right column: The same heat map without the deconvolved events. (c) Left: Cumulative histogram of the shuffle-subtracted sparsity scores from all the shared neurons on Day 1 and Day 6 (n=10 mice; *** P≤ 0.001 Kolmogorov-Smirnov test). Right: The average shuffle-subtracted sparsity scores of shared neurons on Day 1 and Day 6 (n= 10 mice; ** P≤ 0.01; paired *t*-test). (d) Left: Cumulative histogram of the shuffle-subtracted sparsity scores from all neurons imaged on Day 1 and Day 6 (n=10 mice; *** P≤ 0.001; Kolmogorov-Smirnov test). Right: The average shuffle-subtracted sparsity scores of all neurons imaged on Day 1 and Day 6 (n= 10 mice; * P≤ 0.05; paired *t*-test). (e) Decoding the location of the animal from the CA1 population activity using a neural network supervised deep learning approach (blue and red rectangles represent adjacent bins or chunks of the neural data inputted into the decoder). (f) Example trial of location decoding of x and y coordinates in the maze in separate graphs on Day 1 and Day 6 as a function of time. Green traces reflect the true position of the animal, and gray traces depict the decoded position. (g) Shuffle-subtracted position decoding error calculated by least mean squares on Day 1 and Day 6 (n= 10 mice; * P≤ 0.05). Right: Shuffle-subtracted correlation coefficient (r) between the decoded and true x,y positions on Day 1 and Day 6 (n= 10 mice; * P≤ 0.05). Statistical analysis performed using a paired *t*-test.

### Improvement of population decoding for position with learning

Since the single-cell analysis suggested improved selectivity for space after spatial learning, we asked whether the CA1 neuronal population allows more accurate decoding of the location of the animal after learning. We trained a feedforward artificial neural network (FFN) (*34*), to decode the location of the animal from the calcium traces of the imaged CA1 neurons before and after spatial learning (Fig. 2, E). Again, we only analyzed a comparable number of neurons that were active both on Day 1 before learning and Day 6 after learning. We also equalized the training and testing time between Day 1 and Day 6 for a fair comparison (see Methods). Examination of multiple example trials demonstrated that the decoding appeared to improve with learning (Fig. 2, F). To quantify improvements in decoding performance with learning, we computed the decoded spatial location error by calculating the distance between the decoded location and the original location of the animal (see Methods). To ensure that changes in the patterns of behavioral exploration did not influence our decoder results, we calculated the shuffled decoded error and subtracted the 95^th^ percentile of the shuffled decoded error from the recorded decoded error for Day 1 and Day 6 accordingly. Indeed, there was a significant reduction in the shuffle-subtracted decoding error for location on Day 6 compared to Day 1, demonstrating that decoding performance improved with learning. The shuffle-subtracted decoded error went below zero on Day 6 as the recorded decoded error was lower than the shuffled decoded error (Fig. 2, G). To further demonstrate that the decoder could track the spatial trajectory of the animal more accurately after learning, we calculated the correlation coefficient between the decoded x and y coordinates and the true x and y coordinates both on Day 1 and Day 6. Again, to account for changes in exploration from Day 1 to Day 6, we shuffle-subtracted the 95^th^ percentile of the measured correlation coefficients. We found that on Day 6 there was a significant rise in the shuffle-subtracted decoded correlation compared to Day 1, demonstrating an improvement in the decoded spatial trajectory of the animal (Fig. 2, G). To account for the behavioral bias, we subsampled Day 1 behavior by Day 6, choosing only locations visited on both days while equalizing recording time on both days. The decoder consistently predicted the location and trajectory of the animal more accurately on Day 6 compared to Day 1 (Supplement Fig. 2, G-H). Together, these results indicate that the location of the animal can be more precisely decoded from the CA1 neuronal population after spatial learning.

### Viral CRISPR-CasG deletion of the neuronal NMDA receptor subunit GluN1 in dorsal hippocampus impairs synaptic plasticity and spatial learning in the MWM

Next, we sought to elucidate the neuronal mechanism that drove the increases we found in spatial selectivity and behavioral performance over the course of learning. It is known that the pharmacological antagonism of NMDA-receptors or transgenic knockout of NMDA receptor subunits in the forebrain impairs synaptic plasticity and spatial learning in the MWM (*35–37*). However, whether dorsal hippocampus NMDA receptor activation was also essential for the improvements in hippocampal spatial coding during spatial learning is not yet known. To answer these questions, we used a novel viral CRISPR-Cas9 strategy to delete *Grin1*, the gene encoding the essential NMDA receptor subunit GluN1, in dorsal hippocampal neurons (*38*, *39*). We confirmed a bilateral dorsal hippocampal injection of AAV virus expressing a small Cas9 (SaCas9) and a guide RNA targeting the *Grin1* gene (GluN1-CRISPR) under the neuronal promoter (*hSyn1*), decreased *Grin1* gene expression in the animals that underwent the imaging experiments. Control animals were injected with an AAV containing SaCas9 without the guide RNA (*40*). One of the dorsal hippocampi was co-injected with an AAV-syn-GCaMP7f for probing the physiological effects of the NMDAR deletion (Fig. 3, A). We allowed 6 weeks for the deletion of the *Grin1* gene (*39*). To characterize *Grin1* deletion, we performed RNA-scope and quantified its transcripts in the dorsal hippocampi. Injection of GluN1-CRISPR reduced *Grin1* mRNA significantly by 60 +/- 15 percent in both dorsal hippocampi (as measured by total expression in CA1, CA3, and dentate gyrus) as well as specifically by 80 +/- 6 percent in CA1 compared to controls (Fig. 3, B-C). We calculated the proportion GCaMP7f neurons co-expressing GluN1-CRISPR (by staining for SaCas9-HA) and found that 62 +/- 11 percent of GCaMP7f expressing neurons also expressed GluN1-CRISPR in CA1 (Fig. 3, D). A similar proportion of GCAMP7f expressing neurons expressed the control viral construct (Fig. 3, D). There was very little expression of the GluN1-CRISPR outside the dorsal hippocampus (Supplement Fig. 3, A-B).

**Fig 3:**
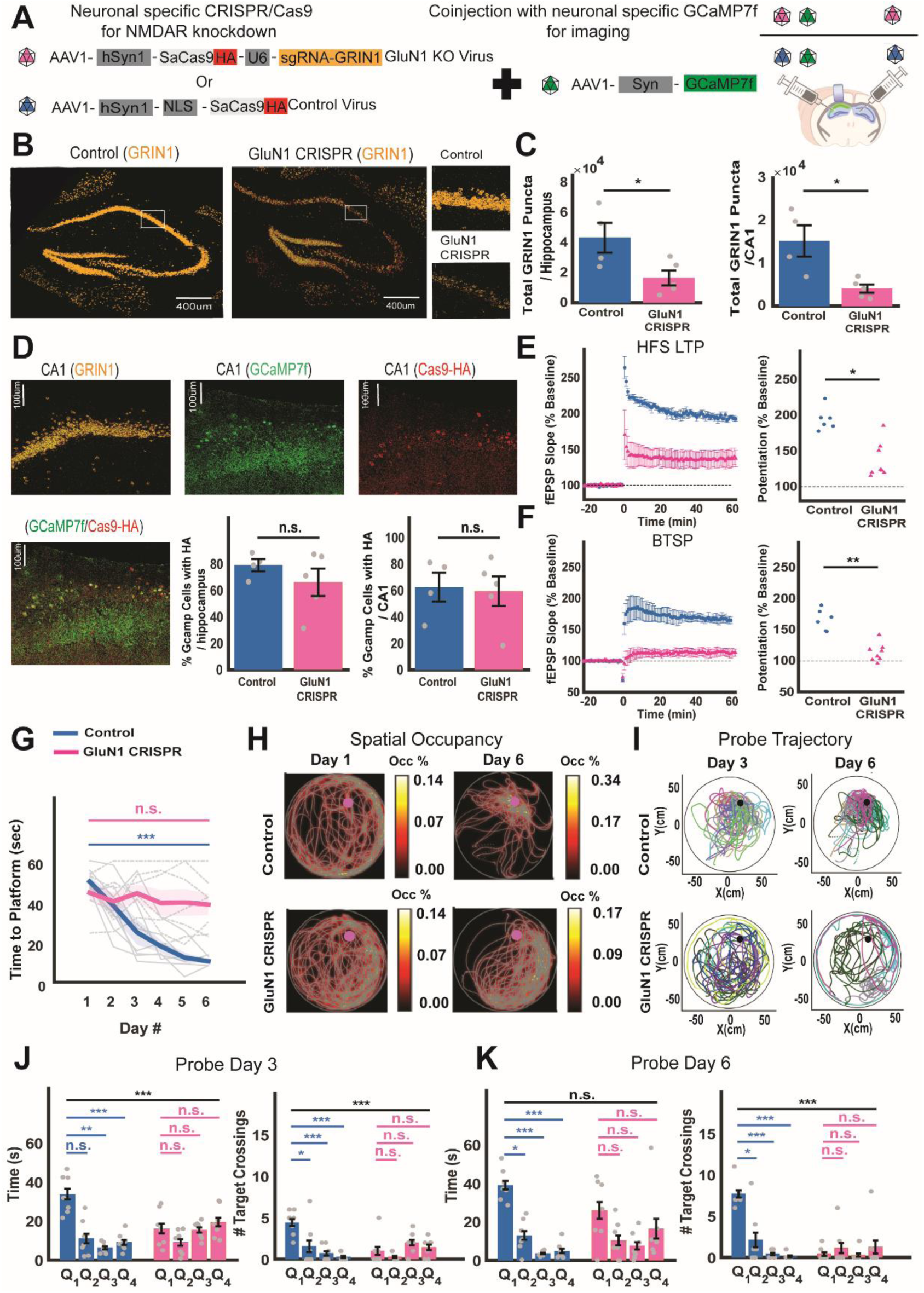
Dorsal hippocampus CRISPR knockdown of neuronal GluN1 reduces burst-dependent synaptic potentiation and spatial learning in the MWM. (a) AAVs used for neuronal-specific CRISPR-Cas9 gene knockdown in bilateral dorsal hippocampi as well as co-injected AAV for expression of GCAMP7f in the right dorsal hippocampus. (b) Confocal images of RNA scope for the *Grin1* mRNA in mice injected with the control virus or the GluN1-CRISPR virus. Insets show magnified images of CA1 in both groups. (c) Quantification of the total amount of *Grin1* stained puncta in control virus-injected and GluN1-CRISPR injected animals, either analyzed across all hippocampal subfields (left; n= 4 controls; n= 5 GluN1-CRISPR; * P≤ 0.05), or just in CA1 (right; n= 4 controls; n=5 GluN1-CRISPR; * P≤ 0.05). (d) Confocal image of an enlarged part of CA1 stained for *Grin1* mRNA in orange, GCamp7f in green and CRISPR-HA tag in red. Co-stained cells expressing HA and GCaMP7f are marked in yellow. Graphs show the number of co-stained neurons in control virus and GluN1-CRISPR injected animals that underwent imaging experiments, either analyzed across all hippocampal subfields (left; n=4 controls; n=5 GluN1-CRISPR; P= 0.336), or just in CA1 (right; n=4 controls; n=5 GluN1-CRISPR ; P= 0.84). (e) Left: HFS induced LTP (100Hz delivered at time = 0) in slices of mice injected with the control virus or the GluN1-CRISPR virus. Right: HFS-induced LTP is significantly reduced in slices of GluN1-CRISPR injected mice (control: 6 slices, n=3 mice; GluN1-CRISPR: 8 slices, n=4 mice; * P< 0.05). (f) Left: BTSP induced by theta-pulse stimulation (5 Hz stimulation for 15 sec delivered at time = 0) in slices of CRISPR-control and GluN1-CRISPR injected animals. Right: BTSP is significantly reduced in slices of GluN1-CRISPR injected mice (control: 6 slices, n=3 mice; GluN1-CRISPR: 8 slices, n=4 mice; *** P≤ 0.001). (g) Time to platform changes across 6 days of training in the MWM in mice injected with the control virus or the GluN1-CRISPR virus. Control injected mice show normal learning progression while GluN1-CRISPR injected mice exhibit an impairment in learning (blue; n=7 controls; *** P≤ 0.001; magenta; n= 7 GluN1-CRISPR; P= 0.673; paired *t*-tests). (h) Spatial occupancy in the maze across all trials of Day 1 and Day 6 of one animal injected with control virus and one injected with the GluN1-CRISPR virus. (i) Overlaid trajectories of all the virus injected control and GluN1-CRISPR injected mice during performance of the probe trial on Day 3 and Day 6. (j) Left: Spatial memory performance in mice injected with the control virus during the probe trial of Day 3. Quantified as the proportion of time spent exploring the learned platform quadrant – Q1 (blue; n= 7 mice; P= 0.017; ** P≤ 0.01; *** P≤ 0.001) or in mice injected with GluN1-CRISPR virus (magenta; n= 7 mice; P= 0.14; P= 0.89; P= 0.6). Right: Number of platform crossings during the Day 3 probe trial in mice injected with the control virus (blue; n= 7 mice; * P≤ 0.0167; *** P≤ 0.001; *** P≤ 0.001), or in mice injected with GluN1-CRISPR virus (magenta; n= 7 mice; P= 0.37; P= 0.23; P= 0.68). Statistical analysis was performed using two-way ANOVA with repeated measures between the groups for occupancy or crossings separately. In each measure within each group we used planned paired *t*-test comparisons with Bonferroni correction for the different quadrants. (k) Left: Spatial memory performance in mice injected with the control virus during the probe trial of Day 6. Quantified as the proportion of time spent exploring the learned platform quadrant – Q1 (blue; n=7 mice; * P≤ 0.0167; *** P≤ 0.01; *** P≤ 0.001) or in GluN1-CRISPR virus injected mice (magenta; n=7 mice; P= 0.097; P= 0.052; P= 0.46). Right: Number of platform crossings during the Day 6 probe trial in mice injected with the control virus (blue; n=7 mice; * P≤ 0.0167; *** P≤ 0.001; *** P≤ 0.001) or in mice injected with GluN1-CRISPR virus (magenta; n=7 mice; P= 0.36; P= 0.36; P= 0.61). Statistical analysis was performed using two-way ANOVA with repeated measures between the groups for occupancy or crossings separately. In each measure within each group we used planned paired *t*-test comparisons with Bonferroni correction for the different quadrants.

To ensure that GluN1 deletion had a robust functional effect on hippocampal plasticity, we performed field potential recordings from hippocampal slices from a separate group of GLuN1-CRISPR or CRISPR injected controls and measured long-term potentiation (LTP) of Schaffer collateral responses. We reasoned that since NMDA receptor signaling is essential for high-frequency stimulation (HFS) LTP induction, then CRISPR deletion of *Grin1* in a sufficient proportion of neurons should decrease the magnitude of LTP in these slices (*35*, *38*). Indeed, the CRISPR induced *Grin1* deletion resulted in a dramatic reduction in HFS induced LTP compared to controls (Fig. 3, E). Behavioral time-scale plasticity (BTSP), a novel form of plasticity that mediates the rapid formation of place cells in CA1 pyramidal neurons in-vivo, is also NMDAR-dependent (*41*). Thus, we also tested the effects of GluN1-CRISPR deletion on a complex-spike burst-dependent form of BTSP (*42*), and found a very strong reduction in BTSP in the GluN1 CRISPR group in comparison to controls (Fig. 3, F). Therefore, GluN1-CRISPR expression dramatically reduces multiple forms of long-term potentiation in CA1.

We examined the effect of *Grin1* deletion on spatial learning in the MWM. These experiments were performed on the same mice that were examined histologically for *Grin1* deletion and co-expression with GCaMP7f. We found that CRISPR control-injected animals showed a progressive decrease in the latency to reach the platform during the 6 days of the MWM behavior. On the other hand, GluN1-CRISPR injected animals did not show a significant decrease in the latency to platform by Day 6 (Fig. 3, G), indicating that spatial learning was impaired in these mice. On Day 1, both the CRISPR-control and GluN1-CRISPR injected animals searched the maze in a uniform manner. On Day 6 controls chose direct paths to the goal or mostly searched near the platform area, while GluN1-CRISPR injected animals still searched the maze in a mostly uniform manner (Fig. 3, H; see Methods). During the probe memory test, CRISPR-control injected mice mostly focused their search around the target quadrant on Day 3, and this search became even more tightly focused on the target quadrant on Day 6. On the other hand, the GluN1-CRISPR injected mice group searched all quadrants equally on Day 3 and did not search the target quadrant more than the other quadrants on Day 6 (Fig. 3, I-G). To ensure that a lack of motivation or motor deficits could not explain the poor performance of GluN1-CRISPR injected animals, we performed 6 blocks of visible platform experiments after the last day of training over two days (*43*). We found no difference between the CRISPR-controls and GluN1-CRISPR groups in the latency to reach the visible platform (Supplement Fig. 3, C). Together, these results show that viral CRISPR-Cas9 deletion of *Grin1* in the dorsal hippocampus can impair spatial learning in the MWM task.

### Hippocampal NMDA receptors are essential for improved spatial selectivity with learning

To examine whether GluN1-CRISPR deletion prevents the improvement of place selectivity in CA1 neurons with spatial learning, we imaged the activity of CA1 neurons in the same animals that underwent bilateral *Grin1* deletion and the CRISPR control animals co-injected with GCaMP7f as they performed the MWM over 6 days. We imaged nearly 700 neurons across 7 animals per group over 6 days (Supplement Fig. 4, A). We focused on neurons that were active on both Day 1 and Day 6 (nearly 315 shared neurons in each group, with no significant difference in group averages) (Supplement Fig. 4, B). In CRISPR-control injected animals, similarly to previous experiments, some neurons showed scattered firing during Day 1, exhibiting low sparsity for space. On Day 6 however, the same neurons showed much more spatially selective firing in different locations in the maze (Fig. 4, A). In contrast, in GluN1-CRISPR injected mice, most neurons showed scattered firing with no clear location firing on Day 1 and on Day 6 (Fig. 4, A). As before, we calculated the sparsity measure and the shuffled sparsity for each neuron in CRISPR-control and GluN1-CRISPR injected mice. Consistent with the previous experiments, the CRISPR-control group cumulative distribution showed a significant shift to higher shuffle-subtracted sparsity values on Day 6 compared to Day 1. On the other hand, there was no increase in the cumulative distribution of shuffle-subtracted sparsity values in the GluN1-CRISPR group on Day 1 compared to Day 6 (Fig. 4, B). Results were similar when shuffle-subtracted sparsity was averaged for each animal and the values from the two groups of animals were compared. In the CRISPR-controls there was a significant increase in shuffle-subtracted sparsity between Day 1 and Day 6, while in the GluN1-CRISPR group there was no increase in shuffle-subtracted sparsity (Fig. 4, B). Calcium event frequency was similar in the two groups and between Day 1 to Day 6 (Supplement Fig. 4, C). The swimming speed of CRISPR-control injected mice and GluN1-CRISPR injected mice were not different between Day 1 and Day 6 (Supplement Fig. 4, D). Including all neurons imaged in the analysis instead of only those with activity on both Day 1 and Day 6 resulted in similar findings (Fig. 4, C). We again controlled the behavioral bias by subsampling Day 1 behavior by Day 6. Similarly, the results showed an increase in the shuffle-subtracted sparsity in Day 6 as opposed to Day 1 in the shared cells (Supplement Fig. 4, E-F) as well as in all the cells imaged (Supplement Fig. 4, G-H). Therefore, these results showed that the deletion of *Grin1* prevented increases in spatial sparsity of individual cells with learning.

**Fig 4:**
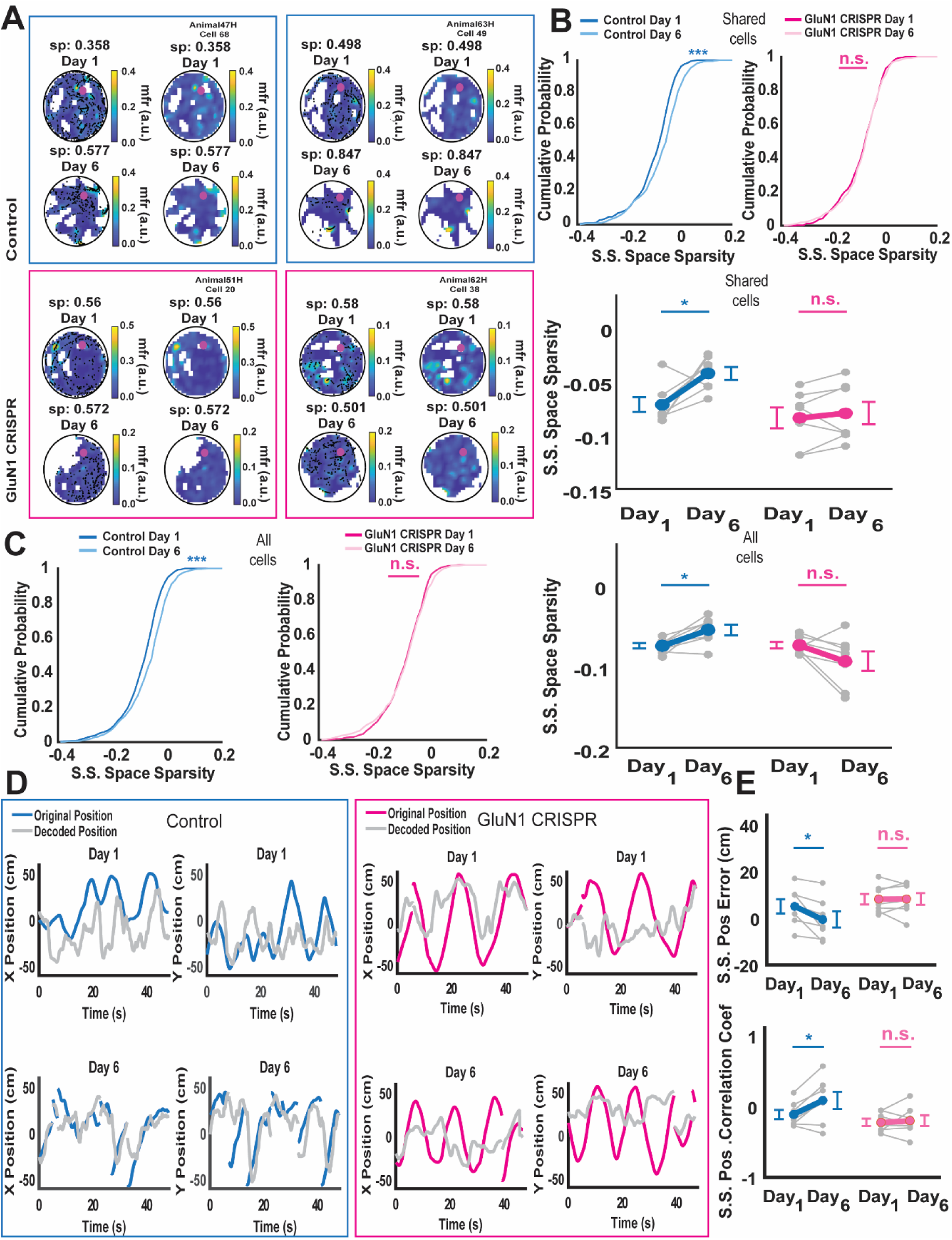
Dorsal hippocampus *Grin1* CRISPR deletion prevents improvement in place selectivity of the CA1 neuronal population after spatial learning in the MWM. (a) Left column: Heat maps of the activity of the imaged neurons on Day 1 and Day 6 with deconvolved events color coded in black. Right column: The same heat maps without the deconvolved events. (b) Left: Cumulative histogram of the shuffle-subtracted sparsity scores from the shared neurons on Day 1 and Day 6 for control virus injected mice (n= 7 mice; *** P≤ 0.001). Right: Cumulative histogram of the shuffle-subtracted sparsity scores from the shared neurons on Day 1 and Day 6 for GluN1-CRISPR injected mice (n= 7 mice; P= 0.927). Statistical analysis was performed using the Kolmogorov-Smirnov test. Bottom: The average shuffle-subtracted sparsity scores of shared neurons on Day 1 and Day 6 of control virus injected mice (blue; n=7 mice; * P≤ 0.05) or GluN1-CRISPR injected mice (magenta; n= 7 mice; P= 0.59). Statistical analysis was performed using paired *t*-tests. (c) Left: Cumulative histogram of the shuffle-subtracted sparsity scores from all the neurons imaged on Day 1 and Day 6 of control virus injected mice (blue; n= 7 mice; *** P≤ 0.001). Middle: The average shuffle-subtracted sparsity scores of all the neurons imaged on Day 1 and Day 6 of GluN1-CRISPR injected mice (magenta; n= 7 mice; P= 0.267). Statistical analysis was performed using the Kolmogorov-Smirnov test. Right: The average shuffle-subtracted sparsity scores of all neurons imaged on Day 1 and Day 6 of control virus injected mice. (blue; n= 7 mice; * P≤ 0.05): or GLuN1-CRISPR injected mice (magenta; n= 7 mice; P= 0.942). Statistical analysis was performed using paired *t*-tests. (d) Left: Example trial of location decoding of x and y coordinates in the maze in separate graphs on Day 1 and Day 6 as a function of time of control virus injected mice, with (blue) representing the original position and (gray) representing the decoded position, or of GluN1-CRISPR injected mice with (magenta) representing the original position and (gray) representing the decoded position. (e) Up: Shuffle-subtracted position decoding error average calculated by least mean squares on Day 1 and Day 6 in control virus injected mice (blue; n= 7 mice; * P≤ 0.05) or in GluN1-CRISPR injected mice (magenta; n= 7 mice; P= 0.52). Bottom: Shuffle-subtracted correlation average between the decoded and true x,y position on Day 1 and Day 6 in control virus injected mice. (blue; n= 7 mice; * P≤ 0.05) or in GluN1-CRISPR injected mice (magenta; n= 7 mice; P= 0.35). Statistical analysis performed using paired *t*-tests.

To determine whether *Grin1* deletion prevented the improvement in position decoding from the population activity during learning, we used the FFN model for spatial location decoding as before. Examination of multiple example trials suggested that spatial decoding improved from Day 1 to Day 6 CRISPR-control injected animals but not in GluN1-CRISPR injected animals (Fig. 4, D). Consistent with this, CRISPR-control animals showed a significant reduction in 95th percentile shuffle-subtracted error between the decoded and real position on Day 6 compared to Day 1. However, in the CRISPR-GLuN1 injected group there was no change in the shuffle-subtracted decoding error of position between Day 1 to Day 6 (Fig. 4, E). CRISPR-control injected mice showed a significant increase in the shuffle-subtracted decoded correlation between Day 1 to Day 6, while the GluN1-CRISPR did not show any change in this measure (Fig. 4, E). Controlling for the behavioral bias, by using only positions visited both in Day 1 and Day 6 while equalizing the recording time, resulted in similar findings. In CRISPR-control injected mice the shuffle-subtracted decoded error was reduced on Day 6 as compared to Day 1 and the shuffle-subtracted correlation coefficient increased, suggesting improved trajectory prediction. On the other hand, GluN1-CRISPR injected mice did not show any change in these measures (Supplement Fig. 4, I-J). Therefore, improvements in hippocampal spatial coding and spatial decoding with learning on the MWM rely on NMDA receptor-dependent synaptic plasticity or activity.

## Discussion

We measured spatial representations in the hippocampus throughout learning by imaging the same population of neurons during free allocentric navigation in 2D. We discovered that CA1 place coding became more precise after learning to navigate to a goal, in accordance with the prediction that cognitive map representation acuity increases with learning of a new environment (*44*). The conditional deletion of NMDA receptor subunits in the dorsal hippocampus using CRISPR, which impaired two forms of long-term potentiation in CA1, prevented learning-related changes in place cell precision and spatial navigation learning.

We examined the changes in hippocampal neural dynamics in a well-controlled 2D free navigation task using distal visual cues (*23*). The use of a water maze enabled us to eliminate any confounding factor of odor left on the goal in dry mazes. Also, since the goal was hidden, the animals could only use the distal visual cues and not use any alternative strategy. Using the water maze also allowed us to directly apply our findings to the multitude of studies performed using this paradigm (*24*, *45*, *46*). To make sure that animals used an allocentric strategy to navigate to the reward each trial had a random different starting point from a total of eight locations. Last, by imaging the exact same cells before and after learning, we were able to directly examine how each neuron in the ensemble, and the ensemble as a whole, changes its response properties. We found improvements in place cell precision and improved decoding of spatial location after learning. Our findings were not explained by changes in the behavior of the animals, as we performed shuffled subtraction of our sparsity and decoding measures, as well as further controls where we only probed changes in firing the same locations visited before and after learning.

We found that NMDA receptor deletion prevents learning-related improvements in place cell precision and subsequent 2D navigation learning. This manipulation also impaired two forms of long-term potentiation recorded in CA1, suggesting these plasticity impairments are responsible for the lack of place precision improvements. However, NMDA receptor deletion was limited to the dorsal hippocampus but was not limited to individual subfields within the hippocampus. Therefore, it is possible that diminished plasticity within DG or CA3 also play a role in the failure of learning-related improvements in place field precision in CA1 with learning (*47*). Because our deletions were performed in adult animals, we can be sure that the results we observed were not the result of loss of NMDA signaling on the development of hippocampal circuits.

Our results differ from those of a recent study in rats performing 2D virtual navigation using visual cues, where there was no improvement in CA1 place coding with learning, but the emergence of path integration representations (*19*). These differences could arise because of poor place cell firing in the restricted virtual reality environment, forcing the animals to adopt a path integration strategy (*20*, *48*).

The improvement in place tuning precision that was found to accompany the allocentric learning suggests that improved precision may explain the improved navigational performance. However, the association between these two phenomena does not prove they are causally related. Further experiments will be needed.

Future studies will also probe the effect of hippocampal subregion and cell-type specific manipulations on place cell precision improvement and 2D spatial learning. Unlike other behavioral arenas, our behavioral paradigm did not allow us to examine changes in vector cells, which only fire when the animal is heading toward the goal location (*49*, *50*). Further experiments in other environments will be needed to determine how these responses change with learning and whether NMDA receptor activation is necessary for these learning-related changes.

### Experimental methods

#### Animals

All experimental protocols were approved by the Chancellor’s Animal Research Committee of the University of California, Los Angeles, in accordance with the U.S. National Institutes of Health (NIH) guidelines. For water maze experiments, 10-week-old hybrid male B6129SF1/J1 mice (Jackson Laboratories, 10143) were group housed (2–5 mice per cage) on a 12–hour light–dark cycle for all experiments. Hybrid mice were used because they were able to learn the task more reliably than C57Bl6 mice(*51*). Male mice were used because they are larger than females at the appropriate adult age, and thus were able to swim and perform the task better while implanted with the miniscope, as assayed in preliminary experiments. Blinding was performed during data collection and analysis.

#### Calcium imaging surgeries

For all surgeries, mice were anesthetized with 1.5–2.0% isoflurane and placed into a stereotactic frame (David Kopf Instruments). Lidocaine (2%, Akorn) was applied to the sterilized incision site as an analgesic, and subcutaneous saline injections were administered throughout each surgical procedure to prevent dehydration.

For calcium imaging experiments, all mice underwent two stereotaxic procedures during the same operative session. Mice were anesthetized and unilaterally injected with 500 nl of AAV1.Syn.jGCaMP7f.WPRE (2.2 x 10^13^ GC/ml) virus at 1 nl per second in the dorsal CA1 (-2 mm anteroposterior relative to bregma, +2.0 mm mediolateral from bregma, and 1.6 mm ventral from the skull surface) using a Nanoject microinjector (Drummond Scientific). They then underwent a GRIN lens implantation surgery. A craniotomy 2 mm in diameter was performed above the viral injection site. The cortical tissue above the targeted implant site was carefully aspirated using 27 gauge and 30-gauge blunt needles. Buffered ACSF was constantly applied throughout the aspiration to prevent desiccation of the tissue. The aspiration was ceased after partial removal of the corpus callosum and full termination of bleeding at which point a GRIN lens (1.8 mm diameter, 4.31 mm length, 0.25 pitch, 0.50 numerical aperture; Edmund Optics) was stereotaxically lowered to the targeted implant site (-1.3 mm dorsoventral from the skull surface relative to the most caudal point of the craniotomy). Cyanoacrylate glue and dental cement (Lang Ce) were used to seal and cover the exposed skull, and Kwik-Sil (MPI) covered the exposed GRIN lens. Carprofen (5 mg kg^−1^) and dexamethasone (0.2 mg kg^−1^) were administered during surgery and then daily for 7 days after surgery. Amoxicillin (0.25 mg ml^−1^) was delivered for 7 days in the drinking water. Animals were anesthetized again 3 weeks later, and a miniature microscope locked onto an aluminum baseplate was placed on top of the GRIN lens. The baseplate and miniscope assembly were adjusted until blood vessels and labeled neurons could be imaged. The baseplate was cemented into place at this position and the miniscope was unlocked and detached from the baseplate. A plastic cap was locked into the baseplate to prevent debris from settling on the GRIN lens surface.

CRISPR injection was done using the exact same procedure. The right hemisphere was injected with a cocktail containing 500nl of GCaMP7f and 500nl of AAV1.hSYN1.SaCas9.HA (5.04 x 10^13^ GC/ml; control virus) or AAV1.hSYN1SaCas9.HA.U6.sgRNA-Grin1 (4.85 x 10^13^ GC/ml; GluN1 CRISPR virus), both were diluted 1:1 with saline. The left hemisphere was injected with 500nl of AAV1.hSYN1.SaCas9.HA (control virus) or AAV1.hSYN1SaCas9.HA. U6.sgRNA-Grin1 (GluN1 CRISPR virus) accordingly, both were diluted 1:1 with saline.

#### Custom imaging in water maze

The UCLA wire-free miniaturized microscope (*21*) scope was coated with a polymer (Key Polymer, Tough-Seal^tm^ 21) that insulates its printed circuit boards (PCBs). The compound was applied with an applicator gun (DMA 50; Tough-Seal) on the electrically exposed parts of the PCB. The polymer was viscous at first and then cured into a rubbery form for long-term durability.

3 Mylar spherical helium-filled balloons (18” in diameter) were used to reduce the weight load on the animal’s head to avoid sinking. The balloons were attached to the wire-free scope using cords through a custom metal holder attached to the body of the scope.

During each day of imaging, the miniscope was kept on the head of the animal for all of the day’s imaging sessions. This was done to avoid movements in the imaged field of view. The miniscope was removed at the end of each day.

#### Morris water maze

Imaging in the Morris water maze (MWM) was performed in a circular maze (120 cm in diameter; Maze Engineers) filled with water mixed with nontoxic white paint (Acrylic white paint – Blick). The distal cue navigation maze had a submerged platform (10 cm in diameter) located in one quadrant. The maze had 8 fixed starting points that were chosen randomly on each trial. Animals ∼13 weeks old were imaged in the maze every other day for a total of 6 imaging sessions. We utilized a well-established behavioral protocol of 4 consecutive trials run in 3 blocks to reach a total of 12 trials daily for increased imaging time (*4*). Each day included 3 blocks separated by 90 minutes of 4 back-to-back trials. In each trial, the animal was given 60 seconds to locate the platform. If the mouse failed to find the platform, the trial was terminated at 60 seconds and mice were removed from the water and placed on the platform. Animals were placed on the platform before and between each trial for 10 seconds. Animals were on the start location with their back facing the maze. Distal cues were placed at 3 locations: at East, North and West, when the center of the maze is considered as the reference 0 coordinate. West – black triangle sticker on a white wall (height – 10”; sides-12”) located 18” from the maze. North – black rectangle sticker on a white wall (height - 7”; side-12.5”) located 14” from the maze. East-a black cabinet pole on a white wall background (height – 42”; side -1.5”) located 14” from the maze. South – open to the experimenter.

Animals were given two probe trials at end of the training trials on Day 3 and Day 6. The probe was done 1 hour after the last block of each day. During the probe trials the platform was removed and animals were allowed to search freely for 60 seconds.

After the probe of Day 6, two more blocks of 12 back-to-back trials were added to increase the recording time of the last day. Blocks were separated 90 minutes apart to allow the animals to rest. In trained animals, each trial was relatively short as animals learned to find it at maximal performance. In the CRISPR-GluN1 deletion group, 2 out of 9 animals that exhibited thigmotaxis on Day 1 or on Day 6 (i.e. swimming in circles while touching the walls) and were excluded from further analysis. We excluded these animals because they had insufficient coverage of the maze to allow for proper analysis of spatial representations.

Visible platform tests were done after the experiment. The platform was moved to a new quadrant and a flag was attached, extending 15 cm above the water. Animals were given the same regimen of trials as the hidden platform test for 2 days with 3 blocks in each day consisting of 4 trials back-to-back.

#### Behavioral tracking

We built a stereo camera system to record behavior in the maze from diagonal angles, using monochrome CMOS cameras (Chameleon 3, Flir) with magnifying lenses (Computar). Cameras were positioned above the maze at an angle of 45 degrees from it. One camera was in the southwest corner of the maze at a distance of 14”; the other was at the east corner at 14” distance as well.

An Arduino-generated 20 Hz TTL signal synchronized the cameras and allowed us to record at the same frame rate as the wire-free miniscope. We used SpinView 1.13.0.33 software for controlling the cameras and saving the videos (.AVI).

We processed these videos with custom-written code in MATLAB (Mathworks, Natick, MA) that identified the dark pixels representing the mouse on the background of the white water maze to obtain the x, y coordinates of the mouse from each camera view. We then combined two camera views to obtain the 3D positions. We first calibrated the alignment using a 4 × 6 checkerboard with 4.5 × 4.5 mm square sizes. We then used a Rotation matrix (Euler quaternions) to make the 3D positions flat on the z-axis, as the 3D coordinates calculated by the calibration algorithm are perpendicular to the diagonal axis of the cameras. Thus, we created the trajectory of the animal as if it was recorded from above. We normalized the x- and y-axes to reach the exact 60 cm radius of the maze.

#### Calcium imaging analysis

Wire-free miniscope data was extracted from microSD cards and saved as uncompressed 8-bit AVI files for processing and analysis. For each day, we aligned and concatenated the movies in the MWM training blocks into one video file. NoRMCorre algorithm with non-rigid registration was used to correct for frame-to-frame translational shifts of the brain during animal movement (*52*). Constrained non-negative matrix factorization for endoscopic recordings (CNMF-E) was used to identify and extract the spatial shapes and fluorescent calcium activity of individual neurons (*53*). Fast online deconvolution for calcium imaging (OASIS toolbox) was used to deconvolve the fluorescent activity extracted from each neuron with an AR1 constrained model (*54*). The resulting measure can be interpreted as the probability of a neuron being active at each frame, scaled by an unknown constant. We thus referred to this measure in arbitrary units. Raw calcium traces were used for neural network decoding.

#### Tracking the same cells across days

To track the same neurons across multiple days of calcium imaging, we applied the CellReg (*27*) package (https://github.com/zivlab/CellReg). The spatial footprints of neurons from each day of recording were entered as inputs to CellReg GUI, which then computed a probabilistic model that the spatial footprints were from the same cell across days using spatial correlations and centroid distances of the spatial footprints. For each animal, a probability distribution of the nearest neighbors suspected to be same cells across days was plotted according to the centroid distance and spatial correlation grade. Similarly, another distribution of second nearest neighbor cells that are not suspected to be the same cells was plotted using these two measures. The cutoff of the maximal centroid distance and lowest spatial correlation grade to consider it the same cell was always put at the intersection of these two distributions to reduce false positive cells. These numbers varied between animals. No animals with considerable overlap between the two distributions were used. The mapping of all registered cells to their indices in each day was obtained from CellReg in a matrix of size N (registered cells) x M (imaging days). This matrix was used to track the activity of the same cells across different recording days.

#### Molecular cloning for CRISPR/Cas9

To generate pAAV-hSyn1-NLS-SaCas9-HA, a pAAV-gfaABC1D-NLS-SaCas9-HA plasmid was obtained from Addgene (ID #178960; (*39*)) and the gfaABC1D promotor was replaced by the hSyn1 promotor amplified from pAAV-hSyn1-mCherry (Addgene #114472). The resulting plasmid hSyn1-NLS-SaCas9-HA was used as a control. To knockdown the *Grin1* gene, U6-sgGrin1 were amplified from pAAV-FLEX-SaCas9-U6-sgGrin1 (Addgene #124852; (*40*)) and inserted into pAAV-hSyn1-NLS-SaCas9-HA to generate pAAV-hSyn1-NLS-SaCas9-HA-U6-sgGrin1. Both plasmids, hSyn1-NLS-SaCas9-HA and hSyn1-NLS-SaCas9-HA-U6-sgRNA-Grin1 have been deposited at Addgene (Addgene #239970 and #239971 respectively). Molecular cloning was performed using In-Fusion cloning kits (Takara Bio). The fully sequenced plasmids were sent to Vigene Biosciences to generate AAV1 serotypes for each construct yielding a concentration >1.0 x 10^13^ genome copies/ml (gc/mL). The cloning and sequencing strategies were designed with the SnapGene software (v7.0.3, Insightful Science).

#### Immunohistochemistry (IHC) and in situ hybridization (ISH)

Mice were deeply anesthetized with isoflurane and transcardially perfused with phosphate-buffered saline (PBS), followed by 4% paraformaldehyde (PFA). Brains were quickly removed and fixed in 4% PFA for 24 hours at 4 °C before paraffin embedding. The formalin-fixed paraffin-embedded samples were cut into 5 µm sections for staining. In situ hybridization was performed with the RNAscope Multiplex Fluorescent Reagent Kit v2 (ACD #323100). Probes against *Grin1* mRNA (ACD # 431611) were applied according to the manufacturer’s protocol with the following modifications: epitope retrieval was performed for 10 minutes and Protease Plus treatment was performed for 5 minutes, and the TSA Vivid fluorophore 650 (ACD #323273) was applied at 1:3000. IHC was performed following the completion of the RNAscope signal amplification steps, as previously described (*55*). Sections were incubated at room temperature in primary antibodies (anti-GFP 1:1000 Abcam # ab183734, RRID:AB_2732027; anti-HA 1:500 Novus Cat# NB600-362, RRID:AB_10124937) for 2 hours followed by secondary antibodies (Thermo Fisher Scientific # A-21206, RRID:AB_2535792; Thermo Fisher Scientific Cat# A-21432, RRID:AB_2535853) for 1 hour. Nuclei were counterstained with DAPI. The sections were mounted with Prolong Gold and allowed to dry overnight prior to imaging.

#### Confocal imaging

Fluorescent imaging was performed on a Zeiss LSM 800 confocal microscope equipped with an inverted Axio Observer.Z1 stand and a Plan-Apochromat 10x/0.45 air objective (image to show extent of HA signal) and a 40x/1.4 oil immersion objective (image for quantification). Fluorophores included DAPI (excitation 353 nm, emission 465 nm), Alexa Fluor 488 (493/517 nm), Cy3 (548/561 nm), and Alexa Fluor 647 (653/668 nm), with a pinhole set to 1 Airy unit for the longest wavelength. Detection was carried out using GaAsP photomultiplier tube detectors. Images were acquired at a resolution of 512×512 pixels. Imaging and tile stitching were performed using Zeiss ZEN software.

#### Quantification of IHC-ISH

QuPath software (v0.5.1) was used for quantification (*56*). Cell segmentation was based on the DAPI counterstain. For quantification of *Grin1* mRNA puncta, the “subcellular spot detection” function was applied with an expected spot size of 1 µm^2^ (range 0.5-2 µm^2^). GCaMP-positive cells were classified using a threshold based on the mean cellular anti-GFP signal. HA-positive cells were classified using a threshold based on the sum of the nuclear anti-HA signal.

#### Slice preparation for electrophysiology

Hippocampal slices from the dorsal third of the hippocampus were obtained from 2- to 3-month-old, male B6129SF1/J hybrid mice injected with AAV1-hsyn1-SaCas9-sgRNA-Grin1 and their AAV1-hsyn1-SaCas9 injected controls. Mice were deeply anesthetized with isoflurane and, following cervical dislocation, the brain was removed and placed in cold (∼4°C), oxygenated (95% O_2_/5% CO_2_) ACSF containing 124 mM NaCl, 4 mM KCl, 25 mM NaHCO_3_, 1mM NaH_2_PO_4_, 2 mM CaCl_2_, 1.2 mM MgSO_4_, and 10 mM glucose (all obtained from Millipore-Sigma). Both hippocampi were then dissected from the brain, and a manual tissue slicer was used to prepare 400-µm-thick slices. The CA3 region was removed, and slices were transferred to interface-type chambers continuously perfused with ACSF (2-3 ml/min) and allowed to recover (at 30°C) for at least 2 hours before recordings.

Electrophysiological slice recordings: Extracellular recordings were done using slices maintained in interface-type recording chambers perfused (2-3 ml/min) with ACSF. A bipolar, stimulating electrode fabricated from twisted strands of Formvar-insulated nickel-chromium wire (A-M Systems) was placed in stratum radiatum to activate Schaffer collateral/commissural fiber synapses onto CA1 pyramidal cells (basal stimulation rate = 0.02 Hz). Field EPSPs (fEPSPs) were recorded in stratum radiatum using low-resistance (»10 MW) glass microelectrodes filled with ACSF and a MultiClamp 700B amplifier (Molecular Devices). Signals were low-pass filtered with a cutoff frequency of 2 kHz and digitized at 10 kHz. After determining the maximal amplitude of fEPSPs evoked by presynaptic fiber stimulation, the stimulation intensity was adjusted to evoke fEPSPs with an amplitude ∼50% of the maximal amplitude. fEPSPs (evoked throughout an experiment at 0.02 Hz) were collected and analyzed using pClamp10 software (Molecular Devices). Conventional Hebbian LTP was induced using high-frequency synaptic stimulation (two, 1-second-long trains of 100 Hz synaptic stimulation delivered with an inter-train interval = 20s). A complex-spike burst-dependent form of BTSP (*42*) was induced using theta-pulse stimulation (single pulses of presynaptic fiber stimulation delivered at 5 Hz, duration = 15s). Two slices/protocol were recorded using slices obtained from the same animal and the results were averaged. Average slopes of fEPSPs (normalized to baseline) were recorded 55-60 minutes after LTP induction were used for statistical comparisons. Statistical tests (Student’s *t*-tests with n = the number of animals/group) were performed using SigmaPlot 12.5 (Grafiti).

### Calcium imaging analysis

#### Spatial tuning analysis

The circular maze was discretized into spatial bins each measuring 4 × 4 cm. For each bin, neuronal firing activity was estimated as the sum of the deconvolved time-varying calcium event activities divided by the total time the animal occupied that bin while moving at speeds greater than 3 cm/s. Spatial bins with an occupancy duration of less than 0.025 s were excluded from further analysis.

#### Reducing the bias when comparing neural tuning selectivity across days

To reduce the bias of spatial tuning selectivity between day 1 and day 6, the total duration of data used for spatial rate map construction was equalized across sessions. For each neuron, trials were randomly subsampled from both day 1 and day 6 such that the total amount of data was matched between sessions. This subsampling procedure was repeated 10 times.

#### Measure of selectivity for the spatial rate map

One commonly used metric for quantifying the degree of selectivity in a rate map is the sparsity (*S*P), which is defined as follows:

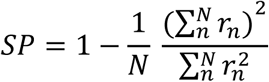

where *N* represents the total number of bins or conditions, such as spatial bins, and *r_n_* denotes the firing rate in the n-th bin. The sparsity is a dimensionless metric, providing insight into the selectivity of a neuron’s firing across various conditions, such as positions. The sparsity ranges from 0 to 1, with a higher value indicating greater selectivity of firing in specific bins (*29*).

#### Normalization of spatial sparsity using shuffled controls

To further reduce the behavioral bias in the comparison of spatial tuning selectivity between day 1 and day 6, sparsity values were normalized relative to shuffled controls. Shuffled datasets were generated by circularly shifting the animal’s position trace by a random temporal offset, thereby preserving the temporal structure of neural activity while disrupting its relationship to spatial position. This shuffling procedure was repeated 50 times for each neuron. For each shuffle, spatial rate maps were recomputed and sparsity was calculated, yielding a distribution of shuffled sparsity values. The 95th percentile of this shuffled sparsity distribution was then subtracted from the original (unshuffled) sparsity value to obtain a normalized sparsity measure for each neuron. This normalization procedure was repeated 10 times, and normalized sparsity values were averaged across repetitions.

[umaht2]

### Behavior-matched control analysis for spatial tuning selectivity

To control for potential differences in behavioral sampling between Day 1 and Day 6, we performed an additional behavior-matched control analysis. Specifically, spatial analyses were restricted to locations that were visited on both Day 1 and Day 6 for each animal. Only spatial bins that met this criterion were included in the construction of spatial rate maps, thereby ensuring that spatial tuning comparisons were based on almost-identical regions of the environment across sessions.

In addition, to prevent bias arising from unequal sampling duration, the total amount of data used to compute spatial sparsity was equalized between Day 1 and Day 6. For each neuron, data from the overlapping spatial bins were randomly subsampled such that the total duration of included data was matched across the two days. Spatial sparsity was then computed using this behavior-matched and duration-matched dataset. This control analysis ensured that observed differences in spatial tuning selectivity were not driven by differences in spatial coverage or behavioral sampling between sessions.

### Feedforward neural network–based decoding

To decode the animal’s position, we implemented feedforward neural networks (FFNs). Neural and behavioral data were first preprocessed and temporally aligned prior to decoder training.

### Data preprocessing and feature construction

Calcium fluorescence signals (ΔF/F) from each neuron were discretized into non-overlapping 100 ms temporal bins, and the average fluorescence value within each bin was computed. Behavioral variable, i.e., position, was similarly discretized into 100 ms bins and averaged within each bin to ensure temporal alignment with the neural data.

To decode the behavioral variable at the *i*-th time bin, we constructed neural feature vectors that incorporated temporal context by including 2*k* + 1consecutive bins of neural activity for each neuron: *k*bins preceding the *i*-th bin, the concurrent *i*-th bin, and *k*bins following it. Unless otherwise specified, *k* = 10, corresponding to a temporal window of 2.1 s centered on the decoded time point.

Feature matrices were generated by concatenating these temporally extended neural signals across all recorded neurons. Each feature matrix consisted of *M*rows, corresponding to the number of time points used for training, and *N* × (2*k* + 1)columns, where *N*denotes the number of neurons. For each neuron, the calcium fluorescence values from the 2*k* + 1bins were extracted and concatenated, and this process was repeated across all neurons to form the full input vector for the FFN.

### FFN architectures for decoding behavioral variables

The input layer consisted of *N* × (2*k* + 1) nodes. For position decoding, we employed a feedforward neural network with two hidden layers, each containing the same number of nodes as the input layer. A dropout rate of 0.2 was applied during training, and the network was trained for 15 epochs. The output layer comprised two nodes representing the decoded x- and y-coordinates of the animal’s position at the *i*-th time bin.

### Decoding performance metrics

To quantify decoding performance and assess improvements associated with learning in the Morris water maze task, we computed the mean positional decoding error as:

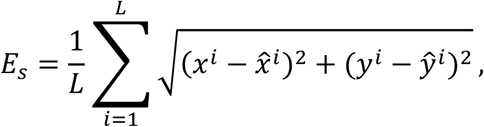

where *x^i^* and *y^i^* denote the true x- and y-coordinates at the *i*-th time bin, *x̅^i^* and *y̅^i^* represent the corresponding decoded coordinates, and *L* is the total number of decoded time bins.

As an additional metric for evaluating position decoding performance, we computed the two-dimensional correlation between the true coordinate matrix *A* = [*X*, *Y*]and the decoded coordinate matrix *B* = [*X̅*, *Y̅*], defined as:

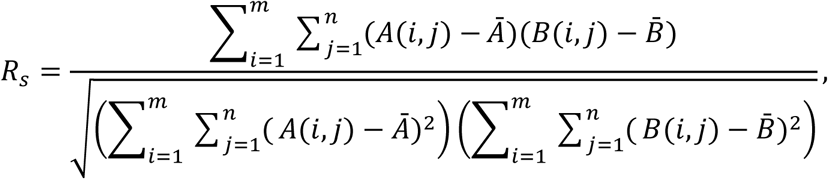

where *m* and *n* denote the number of rows and columns of the coordinate matrices, respectively, *A*(*i*, *j*) and *B*(*i*, *j*) represent individual matrix elements, and *A*^ˉ^ and *B*^ˉ^ are the mean values of matrices *A* and *B*.

### Reducing the bias when comparing decoding performance across days

To reduce bias in the comparison of decoding performance metrics between Day 1 and Day 6, the total amount of data used for decoding was equalized across sessions. For each animal, trials were randomly selected from Day 1 and Day 6 such that the total duration of data was matched between the two days. Following data-length equalization, the dataset from each day was randomly partitioned into training and testing subsets, with 80% of the data used for training and the remaining 20% reserved for testing. This procedure ensured that differences in decoding performance between Day 1 and Day 6 were not driven by unequal data availability or trial selection.

### Normalization of decoding performance using shuffled controls

To further reduce the behavioral bias of decoding performance metrics between Day 1 and Day 6, we normalize decoding performance metrics, we generated shuffled control datasets by circularly shifting the calcium fluorescence trace of each neuron by a random temporal offset. This procedure preserved the temporal structure of neural activity while disrupting its alignment with behavioral variables. The shuffled neural signals were then used as inputs to the same feedforward neural network architectures, and decoding performance metrics—including positional decoding error and two-dimensional correlation coefficient—were computed for each shuffle.

This shuffling procedure was repeated 50 times, yielding a distribution of decoding performance metrics. The 95th percentile of the shuffled distribution was then subtracted from the corresponding original (unshuffled) decoding metric to obtain a normalized measure of decoding performance. To further reduce variability due to random subsampling, the entire shuffling and normalization procedure was repeated 10 times, and normalized decoding performance metrics were averaged across repetitions.

*E*_shuffle_-subtracted decoding error = − *p*(*E*_shuffled_, 95)

### Behavior-matched control analysis for decoding performance

To control for potential effects of behavioral differences between Day 1 and Day 6 on decoding performance, we performed an additional behavior-matched control analysis analogous to that used for spatial sparsity. Specifically, decoding analyses were restricted to spatial positions that were visited on both Day 1 and Day 6 for each animal. Only time bins corresponding to these overlapping positions were included, thereby ensuring that decoding performance was evaluated on almost-identical behavioral states across sessions.

In addition, to eliminate bias arising from unequal sampling duration, the total amount of data used for decoding was equalized between Day 1 and Day 6. For each session, time bins were randomly subsampled from the overlapping spatial positions such that the total data duration was matched across days.

### Shuffled normalization of behavior-matched decoding performance

Following behavior- and duration-matching, decoding performance was normalized using shuffled controls. Shuffled datasets were generated by circularly shifting the calcium fluorescence trace of each neuron by a random temporal offset. This shuffling procedure was repeated 50 times, yielding a distribution of decoding performance metrics. The 95th percentile of this shuffled distribution was then subtracted from the original (unshuffled) decoding metric to obtain a normalized measure of decoding performance.

### Statistical analysis

All statistical analysis was conducted using Prism (GraphPad) or MATLAB (MathWorks). Statistical tests used in this study include paired *t*-tests for two samples (as appropriate). For probe analysis with four samples, we used planned paired *t*-tests with Bonferroni correction (* P≤ 0.0167). Two-way ANOVA with repeated measures was used for statistical analysis of all other samples. The Kolmogorov-Smirnov test was applied to cumulative distributions. The significance threshold was held at α = 0.05; Not significant, P> 0.05; *P≤ 0.05; **P≤ 0.01; ***P≤ 0.001. Sample sizes were not predetermined using statistical methods.

## Acknowledgements

We thank A. Arac, T.J. O’Dell, M. Ollivier and L. Sheintuch for feedback on the manuscript. We also thank J. Liu for technical support. This work was supported by the following funding sources: 4R01AG03622, P50HD103557, R01MH132736 and the NSF Mentor award for M.S.

## Ethics declaration

The authors declare no competing interests.

## Data availability

Data and analysis code will be made available in supplementary information and on GitHub after manuscript acceptance. Any additional information to analyze the data reported in this paper is available from the corresponding authors.

